# Future climate and land use change will equally impact global terrestrial vertebrate diversity

**DOI:** 10.1101/2024.12.13.627895

**Authors:** Chantal Hari, Thomas Hickler, Christian Hof, Christopher P.O. Reyer, Inne Vanderkelen, Alke Voskamp, Matthias F. Biber, Markus Fischer, Édouard L. Davin

**Affiliations:** Wyss Academy for Nature at the University of Bern, University of Bern, Bern, Switzerland; Climate and Environmental Physics, Physics Institute, University of Bern, Bern, Switzerland; Oeschger Centre for Climate Change Research, University of Bern, Bern, Switzerland; Senckenberg Biodiversity and Climate Research Centre (SBiK-F), Frankfurt am Main, Germany; Department of Global Change Ecology, Biocenter, University of Würzburg, Germany; Potsdam Institute for Climate Impact Research, Member of the Leibniz Association, Potsdam, Germany; Department of Earth and Environmental Sciences, KU Leuven, Leuven, Belgium; Royal Meteorological Institute Belgium (RMIB), Brussels, Belgium; Senckenberg Research Institute and Nature Museum, Terrestrial Zoology, Frankfurt am Main, Germany; Institute of Plant Sciences, Plant Ecology, University of Bern, Bern, Switzerland

**Keywords:** climate scenarios, shared socioeconomic pathways, biodiversity, species distribution model, species richness, ISIMIP scenario analysis

## Abstract

**Aim:** Terrestrial biodiversity is impacted by both climate and land use change. Yet, future biodiversity projections have rarely considered these two drivers in combination. In this study, we aim to assess the individual and combined impact of future climate and land use change on global terrestrial vertebrate diversity under a “sustainability” (SSP1-RCP2.6) and an “inequality” (SSP4-RCP6.0) scenario.

**Location:** Global land, excluding Antarctica.

**Time period:** 1995, 2080

**Major taxa studied:** Amphibians, birds, and mammals.

**Methods:** We combined global climate-driven species distribution model (SDM) projections of 13903 vertebrates (amphibians, birds, and mammals) with future and present land use projections from the Land Use Harmonization 2 (LUH2) project. We refined the SDM outputs by the habitat requirements of each species using a land use filtering approach. We then analyzed future species richness changes globally, per region, and per land use category, and looked at taxon-specific effects.

**Results:** Under both scenarios, decreases in future species richness dominate at low and mid-latitudes, with climate and land use change playing an equally important role. Land use change can be either an alleviating (SSP1-RCP2.6) or an exacerbating (SSP4-RCP6.0) factor of climate-induced biodiversity loss. Sub-Saharan Africa is projected to become a high-risk area for future land use-driven biodiversity loss under the SSP4-RCP6.0. Under SSP1-RCP2.6, forested and non-forested land areas increase, while SSP4-RCP6.0 leads to higher rates of deforestation and pasture expansion. Mammals experience the largest climate-driven losses, affecting 56.4% of land area under SSP4-RCP6.0, while amphibians are particularly vulnerable to land use-driven losses, especially under SSP4-RCP6.0.

**Main conclusions**: Our results suggest that both climate and land use pressures on biodiversity will be highest in lower latitudes, which harbor the highest levels of biodiversity.

## 1. Introduction

Biodiversity loss, land degradation, and climate change are some of the key environmental challenges faced by humanity (IPBES, 2019; Rosenzweig et al., 2008). The current biodiversity crisis has been referred to as the “sixth mass extinction” in Earth’s history (Bellard et al., 2012). An increasing amount of land has been converted from its natural state to agricultural land, affecting nearly a third (32%) of the global land area over the past six decades, a rate around four times greater than previously estimated from long-term land change assessments (Winkler et al., 2021). This widespread conversion has led to fragmentation, degradation, and the destruction of species habitat (IPBES, 2019). While past biodiversity loss has mainly been driven by changes in habitat due to land use and land cover changes (Sala et al., 2000; Warren et al., 2018), climate change emerges as an additional risk for biodiversity at the global scale (IPBES, 2019; Pecl et al., 2017). Without stringent climate change mitigation, climate change can lead to potentially large contractions in species’ ranges as they may no longer be able to adapt to the environmental conditions in a given region (Bellard et al., 2012; Jenkins et al., 2021; Warren et al., 2013). Moreover, climate and land use change can have interacting effects on species and ecosystems (Brook et al., 2008; Hof et al., 2011; Mantyka-pringle et al., 2012). For instance, land use change can limit species’ abilities to respond to climate change through dispersal (Hof, 2021; Jenkins et al., 2021; Opdam & Wascher, 2004).

To comprehensively understand the intertwined impacts of climate and land use change and their relative importance, it is essential to consider both factors in future biodiversity projections. However, most global research focused solely on the impacts of climate (Biber et al., 2023; Warren et al., 2018) or land use change (Leclère et al., 2020; Newbold et al., 2015; Powers & Jetz, 2019), even though land use change will occur alongside a changing climate (Jenkins et al., 2021; Methorst et al., 2017), with potentially different outcomes. To date, few studies have made first steps to overcome these deficiencies, for example, by combining species distribution models (SDMs) with land use impacts from a statistical model (Newbold, 2018; Newbold et al., 2020; Pereira et al., 2024). Other global studies consider both drivers but do not assess their separate and combined effects (Beyer & Manica, 2020). Hof et al. (2018) evaluated the potential future impacts of climate and land use changes on global species richness of terrestrial vertebrates under a low and high-emission scenario, with large-scale deployment of Bioenergy with Carbon Capture and Storage (BECCS) for climate mitigation (Hof et al., 2018). However, their analysis did not explicitly account for the habitat requirements of individual species, but rather only considered a spatial overlap of projected species distributions with future land use.

In this study, we address the limitations of previous assessments by explicitly incorporating species-specific habitat requirements into SDMs. This allows for a more realistic representation of species’ responses to land use and climate change. Moreover, we integrate the latest land use projections from Integrated Assessment Models (IAMs), combining these with climate-driven SDM outputs to quantify the separate and combined impacts of climate and land use change on biodiversity. Unlike Hof et al. (2018), who focused primarily on the implications of BECCS-related land use change, our analysis includes a broader range of possible futures by considering two contrasting combinations of Shared Socioeconomic Pathways (SSPs) and Representative Concentration Pathways (RCPs): a “sustainability” scenario (SSP1–RCP2.6) and an “inequality” scenario (SSP4–RCP6.0). We assess projected species richness changes globally and regionally across three terrestrial taxa - amphibians, birds, and mammals - covering 13,908 species and combining climate-driven SDM projections with a land use filtering approach that accounts for species-specific habitat preferences. This integrated framework provides a more nuanced understanding of how biodiversity is likely to be affected under different climate and socioeconomic futures.

## 2. Methods

### 2.1. Species distribution modeling

Future projections with climate-driven SDMs for 15496 terrestrial vertebrate species (2964 amphibians, 8493 birds, and 4039 mammals) were obtained from Hof et al. (2018) as probabilities of occurrence for each species. These projections were based on two types of climate-driven SDMs, generalized additive models (GAM) and generalized boosted regression models (GBM). GAM is an additive model, whereas GBM is a classification and regression tree-based modeling approach. These two approaches were selected because they perform comparatively well to other modeling approaches (Araújo et al., 2005; Elith et al., 2010; Hof et al., 2018; Meynard & Quinn, 2007). The SDMs were calibrated using expert range maps from International Union for Conservation of Nature (IUCN) for amphibians and mammals and from Birdlife International and Nature Serve for birds (IUCN, 2023; Lenoir & Svenning, 2015). More methodological details on the climate-driven SDM approach used, including input data, pre-processing, model validation, bioclimatic variable and pseudo-absence selection, as well as spatial autocorrelation, are provided by Hof et al. (2018). The SDMs were driven by bias-corrected climate data from the Intersectoral Impact Model Intercomparison Project (ISIMIP) phase 2b (Frieler et al., 2017, Lange, 2016) based on four global climate models (GCMs; MIROC5, GFDL-ESM2M, HadGEM2-ES, and IPSL-CM5A-LR) from the Coupled Model Intercomparison Project Phase 5 (CMIP5) under a low (RCP2.6) and high warming scenario (RCP6.0). Note that the CMIP5-based climate scenarios are not explicitly tied to a specific SSP, but for simplicity, we refer to both the climate change and the land use change scenarios as “SSP1-RCP2.6” and “SSP4-RCP6.0” since they combine climate information from the RCPs and land use information from the SSPs (see next section).

We based the analysis in this study on a 30-year mean centered around the years 1995, 2050, and 2080. For simplicity, we refer to these periods by their central year, 1995, 2050, and 2080, without repeatedly mentioning the 30-year mean. The results for the projections for 2050 are presented only in the supplementary material (Fig. S1).

### 2.2. Land use filtering

To account for the impact of future land use change on biodiversity, we use historical and future data from the LUH2 dataset (Hurtt et al., 2020). These data come at a 0.25° resolution and were aggregated to 0.5° resolution to match the spatial resolution of the SDM outputs. All LUH2 data before 2015 is based on historical reconstructions and remote sensing (Hurtt et al., 2020). After 2015, data are based on projections from IAMs under different SSPs and RCPs. IAMs provide projections of different modeled futures that combine knowledge on energy use, land use, greenhouse gas emissions, and climate (Riahi et al., 2017).

SSP1-RCP2.6 is based on the IAM IMAGE 3.0 (Stehfest et al., 2014), and SSP4-RCP6.0 is based on the Global Change Assessment Model (GCAM; Wise et al., 2014). SSP1 projects a future under a sustainable development paradigm. It emphasizes high economic growth with environmentally friendly technologies and population growth that decreases in the second half of the 21^st^ century (van Vuuren et al., 2017). SSP4 describes a world with large inequalities in economic opportunity and political power both across and within countries. High-income countries strongly regulate land use change, but in contrast, in poor countries, tropical deforestation continues (Popp et al., 2017). For both scenarios, assumptions about the climate mitigation policies consistent with the respective RCP are added to the SSP baseline scenarios (Hurtt et al., 2020). LUH2 entails 12 land use categories, namely, forested primary land, non-forested primary land, potentially forested secondary land, potentially non-forested secondary land, managed pasture, rangeland, urban land, and five types of croplands. Habitat preferences for each species were derived from the IUCN Habitat Classification Scheme (v.3.1). The 104 IUCN habitat classes were converted using the lookup table from Carlson et al. (2022; Fig. 1) to match the 12 LUH2 land use categories. The 12 LUH2 land use categories were then further grouped into 5 generalized land cover classes (forested land, non-forested land, pasture, cropland, and urban) according to Powers & Jetz (2019). We combined forested primary land and potentially forested secondary land into forested land, all non-forested primary land and potentially non-forested secondary land into non-forested land, managed pasture and rangeland into pasture, and the five crop types into cropland. To support the decision of combining primary and secondary forests into a single ’forested land’ category in our study, we refer to findings by Rozendaal et al. (2019) which demonstrate that secondary forests can rapidly regain species richness, achieving 80% recovery within two decades, yet require centuries to fully match the species composition of old-growth forests. The final 5 land use categories were then linked to each species separately to account for their individual habitat classification. For 260 amphibians, 1233 birds, and 100 mammals we were not able to apply the land use filtering because there was no available information on their habitat preferences or because all grid cells had a land use fraction of zero in areas with non-zero probability of occurrence (this was the case for 16 (of the 260) amphibians, 2 (of the 1233) birds, and 84 (of the 100) mammals). As a result, these species were removed from the analysis, resulting in 13903 species (2704 amphibians, 7260 birds, 3939 mammals) included in our analysis. If for a given species the habitat was labeled as “suitable” according to the IUCN habitat classification, the probability of occurrence projected by the SDM was multiplied by the grid cell fraction occupied by this habitat (Fig. 1). Since a given species can have multiple suitable habitats, the probability of occurrence was weighted by the grid cell fraction occupied by the sum of all suitable habitats (Fig. 1). Since “urban” is not recognized a suitable habitat for any species according to the IUCN habitat classifications, it is excluded from the land use filtering process.

**Fig. 1.**
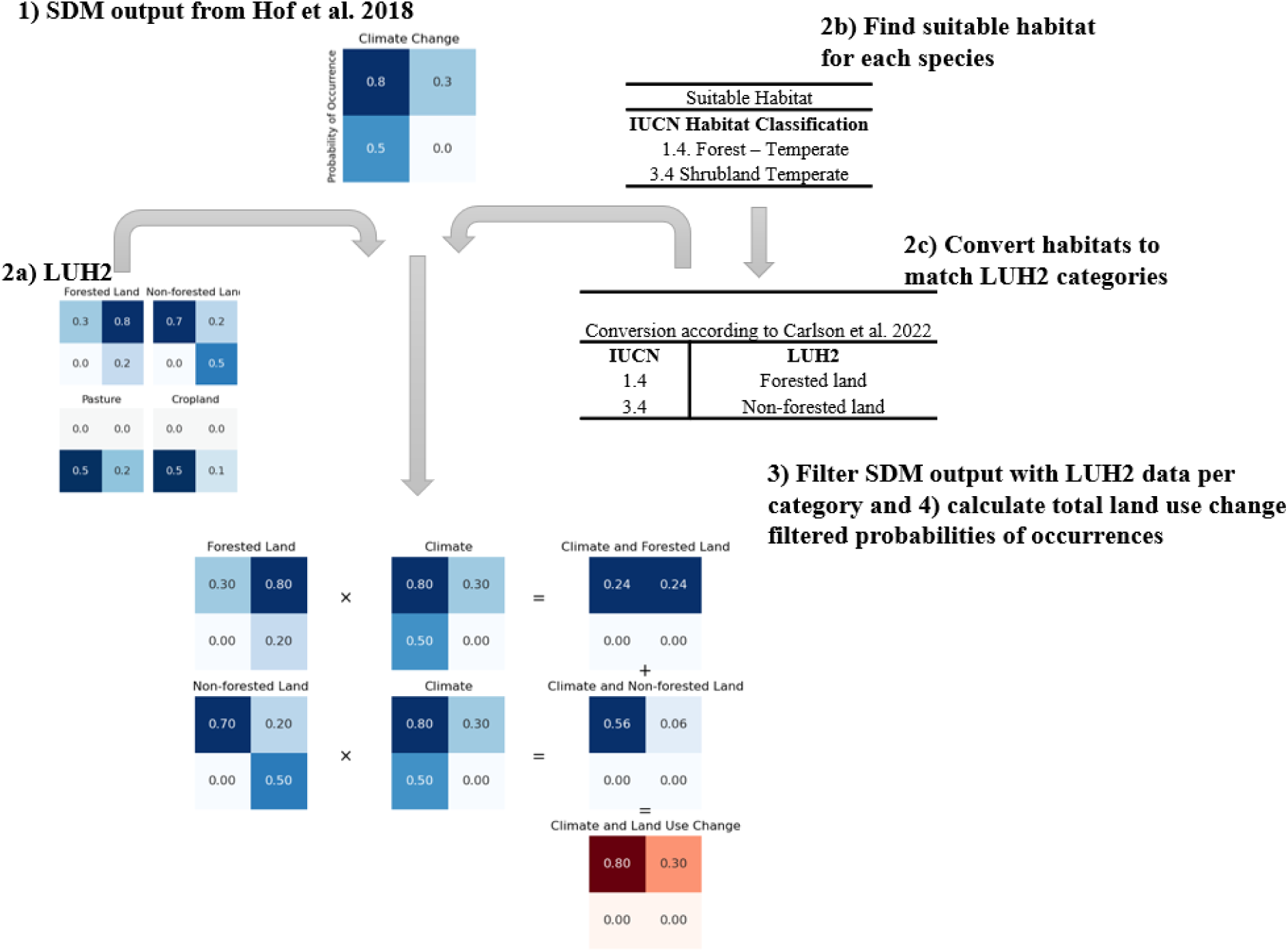
Schematic overview of the different methodological steps of the land use filtering approach. The process is illustrated over 4 grid cells for a single hypothetical species having forested and non-forested land as habitat preferences, with values reflecting changes between future and present-day. 1) We use SDM output from Hof et al., (2018) as probabilities of occurrences. 2) Using the Land Use Harmonization 2 (LUH2) dataset (Hurtt et al., 2020), we classify land into five main categories: forested land, non-forested land, pasture, cropland, and urban areas. Each species’ habitat preference is determined according to the IUCN Habitat Classification Scheme (IUCN, 2023) and matched to the LUH2 land use categories following the conversion table from Carlson et al. (2022). 3) We apply the land use filter by weighing the SDM-derived probabilities of occurrence by the fraction of grid cell occupied by suitable land use categories for each species. 4) These weighted probabilities are then summed to obtain the total land use-filtered probabilities of occurrences.

### 2.3. Calculation of climate-only and land use-only impacts on biodiversity

The combined impact of climate change and land use change is derived by applying the land use filtering approach to the SDM outputs, with present-day land use being applied to the present-day SDM outputs and future land use to the future SDM outputs. The difference between future (2050 or 2080) and present-day (1995) periods is then used to assess the impact on biodiversity. To calculate the effect of climate change only, we also apply the land use filtering approach to the SDM data in order to ensure consistency in the present-day baseline, but land use is kept constant at present-day level. Finally, the impact of land use change was computed by subtracting the climate change-only effect from the combined climate and land use change impact. Given that the land use effect is derived as a residual, it should be noted that it includes potential interaction effects between climate and land use change, arising from the fact that the impact of land use change on biodiversity can be climate-dependent.

### 2.4. Species richness aggregation

For our study, we worked with a large matrix of 13903 species over three taxa, two scenarios (SSP1-RCP2.6 and SSP4-RCP6.0), four GCMs (MIROC5, GFDL-ESM2M, HadGEM2-ES, and IPSL-CM5A-LR), and two SDMs (GAM and GBM). To obtain ensemble projections, we first averaged the SDM outputs and the GCM outputs for each scenario and species. To estimate species richness, the probability of occurrence data per species was first summed across all species for all taxa. scenario, GCM and SDM. To compute regional means, we used region definitions based on the Intergovernmental Platform on Biodiversity and Ecosystem Services (IPBES) assessments (IPBES, 2019). From the 17 IPBES subregions, we derived 10 regions that were used for the analysis (Table S1). The proportion of global land area affected by species richness gain and loss was calculated by summing the area of individual grid cells affected by gain or loss and normalizing the result by the total land area (excluding Antarctica).

### 2.5 Sensitivity analyses

We conducted two sensitivity analyses to test how certain methodological assumptions affect the results. First, because previous studies have shown that different dispersal assumptions represent a major uncertainty in SDM projections, we considered here two dispersal assumptions to test the robustness of our results, following the methods of Hof et al. (2018). The main results presented here are based on the assumption of limited dispersal, in which a buffer of d/4 is applied to each range polygon, with *d* being the diameter of the largest range polygon of a species. In addition, we also present a sensitivity analysis with a “No dispersal” assumption corresponding to a situation where species are not allowed to migrate beyond their current range in the Supplementary Materials (Fig. S3). Secondly, we have tested the sensitivity of the results towards our habitat suitability assumptions. We have applied the land use filter on species with habitat suitability marked as “suitable” according to the IUCN Habitat Classifications Scheme, which is defined as “the species occurs in the habitat regularly or frequently” (IUCN, 2023). For the sensitivity analysis, we have also included species with habitat suitability marked as “marginal” (i.e., “the species occurs in the habitat only irregularly or infrequently” or “only a small proportion of individuals are found in the habitat” (IUCN, 2023).

## 3. Results

### 3.1 Combined species richness changes under climate and land use

To assess future species richness changes, we analyzed the combined and individual impacts of climate and land use change on species richness under two contrasting SSP-RCP scenarios: a sustainability scenario (SSP1-RCP2.6) and an inequality scenario (SSP4-RCP6.0). These scenarios follow distinct land use trajectories (Fig. S8), with SSP1-RCP2.6 assuming strong environmental regulations that reduce tropical deforestation (Hurtt et al., 2020), and SSP4-RCP6.0, assuming limited regulation outside high-income regions, resulting in extensive agricultural expansion at the expense of natural ecosystems. Because species vary in their habitat requirements (Table 1), these land use projections have direct implications for future biodiversity.

**Table 1.** Number of species per taxa and habitat. Species were included if their habitat was categorized as ‘suitable’ according to the IUCN Habitats Classification Scheme. Species can occur in multiple habitats and may thus be listed in multiple categories. Numbers in brackets indicate the relative proportion in reference to the total number of species per taxa.

|  | **Total # of species** | Forested land | Non-forested land | Pasture | Cropland | Single habitat | Multiple habitats |
| --- | --- | --- | --- | --- | --- | --- | --- |
| Amphibians | **2704 (19.5%)** | 2235 (82.7%) | 2211 (81.8%) | 642 (23.7%) | 615 (22.7%) | 660 (24.4%) | 2044 (75.6%) |
| Birds | **7260 (52.2%)** | 5770 (79.5%) | 4644 (64.0%) | 2096 (28.9%) | 1987 (27.4%) | 2622 (36.1%) | 4638 (63.9%) |
| Mammals | **3939 (28.3%)** | 2907 (73.8%) | 2210 (56.1%) | 436 (11.1%) | 542 (13.8%) | 2372 (60.2%) | 1567 (39.8%) |
| **Total** | **13903** | **10912 (78.5%)** | **9065 (65.2%)** | **3174 (22.8%)** | **3144 (22.6%)** | **5654 (40.7%)** | **8249 (59.3%)** |

We find that climate-driven species richness is projected to increase at higher latitudes for both scenarios and widely decreases at mid- to lower latitudes (Fig. 2A & B). Both scenarios show a similar spatial pattern, which is amplified in the SSP4-RCP6.0. Additionally accounting for future land use change leads to more spatial variability of future species richness (Fig. 2E & F). Overall, land use change has a positive impact under SSP1-RCP2.6 (Fig. 2C, while it mostly exacerbates species losses under SSP4-RCP6.0 (Fig. 2D). Global patterns in the climate-driven impact and the combined impact under SSP1-RCP2.6 widely vary worldwide. Sub-Saharan Africa shows the most pronounced negative difference in species richness change between SSP1-RCP2.6 and SSP4-RCP6.0. Especially under SSP4-RCP6.0, species richness mainly increases in the higher latitudes and decreases in the mid-to lower latitudes, also when land use change impact is included (Fig. 2F).

**Fig. 2.**
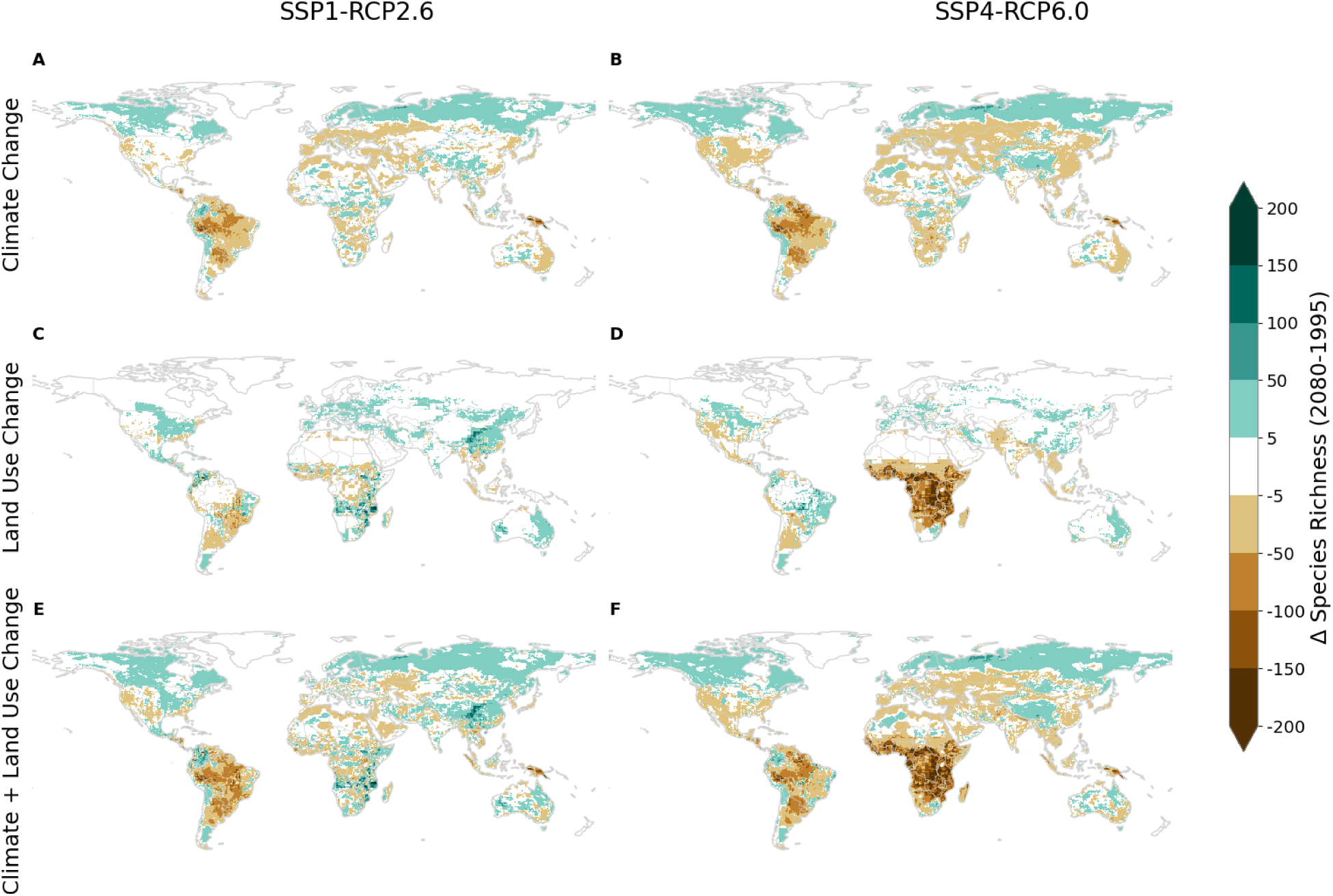
Relative change in species richness due to climate change, land use change, and the combination of both. Projected relative species richness change for the year 2080 compared to 1995 under climate change-only (A and B), land use change-only (C and D), and the combined climate and land use change (E and F). Results are shown for SSP1-RCP2.6 (A, C, and E) and SSP4-RCP6.0 (B, D, and F). Species richness is calculated as the summed probabilities of occurrence over all species of the taxa amphibians, birds, and mammals. The mean over all GCM and SDM combinations is shown here. A minimum threshold of 1e-6 is introduced on the stacked probabilities of occurrences to exclude near-zero values and thus, avoid division by zero for the relative difference calculations. The results for the year 2050 compared to 1995 can be found in the Supplementary Material (Fig. S1) as well as the results for individual taxa (Fig. S4-S6).

The two contrasting land use trajectories directly affect species distribution since habitat preferences differ per species (Table 1). For instance, 78.5% of all species included in this study have forested land as a suitable habitat (i.e., suitable according to the IUCN Habitats Classification Scheme; see Materials and Methods) and will therefore be affected by the decrease in forested land in SSP4-RCP6.0. Conversely, 11.1% of species have pasture as a suitable habitat and thus will be directly affected by the decrease in pasture under SSP1-RCP2.6. Moreover, 56.1% of the species require non-forested land and 13.8% cropland as suitable habitats (Table 1). It is important to note that 59.3% of all included species occur in multiple habitats (Table 1). Thus, species having forest as a suitable habitat are not necessarily forest specialists and can also occur in other habitats.

### 3.2 Species richness changes per region and land use category

Assessing the relative changes in species richness in 2080 compared to 1995 over various world regions (Fig. 2; Fig. 3) shows that climate change alone leads to species richness losses in all world regions except for boreal regions (Eastern Europe and North America) and relative changes are also amplified under SSP4-RCP6.0. Climate-driven species richness loss is highest in South America with an overall relative decrease of -7.2% and -9.1% in SSP1-RCP2.6 and SSP4-RCP6.0, respectively. When combined with land use change, the losses are generally dampened, and gains are amplified in most regions in SSP1-RCP2.6. In SSP4-RCP6.0, however, losses are aggravated and the differences between the two scenarios are exacerbated. In SSP1-RCP2.6, the inclusion of land use change leads to a relative increase in species richness from -1.2% to 2.2% in West, Central, East and South Africa, and higher species richness in most regions except North Africa and Western Asia and South America. The contrary is the case for SSP4-RCP6.0, where the inclusion of land use change leads to a further decrease in species richness in all regions except Central and Western Europe, Eastern Europe, and North America (Fig. 3). This is most pronounced in West, Central, East and South Africa, regardless of the land use category. Overall, the results vary substantially between the different regions and the underlying land use scenarios driven by the respective IAM.

**Fig. 3.**
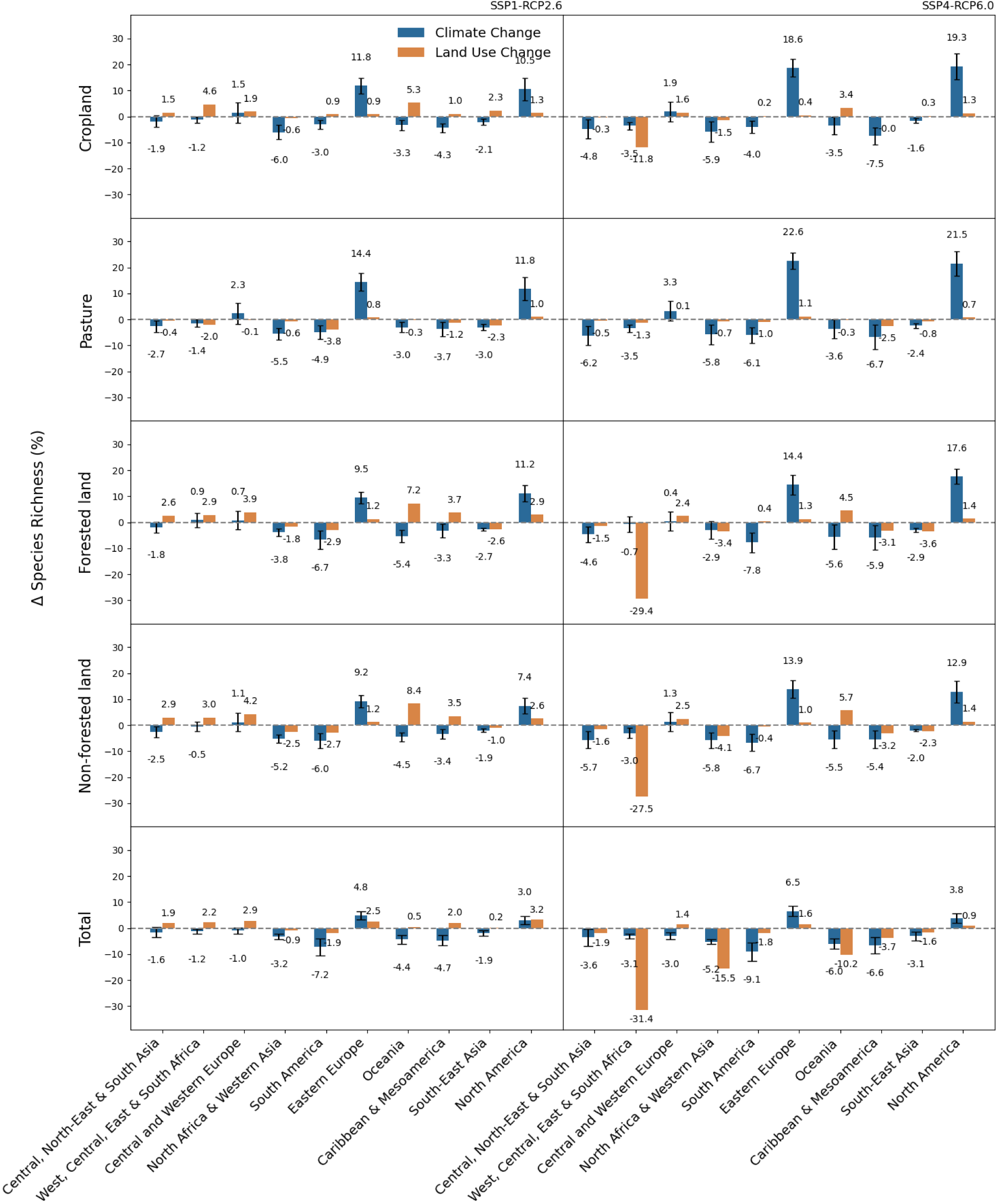
Relative change in species richness from 1995 to 2080 per habitat separately and in total and region. The climate and land use change impact for the two scenarios SSP1-RCP2.6 and SSP4-RCP6.0 per habitat forested land, non-forested land, cropland, pasture, and the total over all habitats. Species richness was calculated separately for each habitat type based on its state at each time point, without tracking transitions between habitat types. The total is based on the relative species richness change from above (Fig. 2). The weighted mean calculation was first done for each combination of GCM and SDM before it was aggregated per region. The land use change-only impact is calculated as the residual between the overall impact and the climate-only impact. The error bars indicate the standard deviation over all GCMs and SDMs. The world regions are based on merged Intergovernmental Science-Policy Platform on Biodiversity and Ecosystem Services (IPBES) regions (Table S1) and results are spatially aggregated using a latitude-weighted mean.

To understand how the different land use narratives translate into biodiversity outcomes, we examined projected changes per land use category. For the three land use categories forested land, non-forested land, and cropland, most of the projected species richness loss is due to land use change for SSP4-RCP6.0. For forested land, -29.4% loss is due to land use change in West, Central, East and South Africa, and only -0.7% loss due to climate change impact, with a standard deviation of 3.4% over all GCMs and SDMs. Similarly, non-forested land projections result in a -27.5% decline in projected species richness loss due to land use change and -3% +/-2.1% SD from climate change in the same region and for the same scenario (Fig. 3). Both forested land and non-forested land are projected to decrease substantially over West, Central, East and South Africa under the SSP4-RCP6.0 scenario (Fig. 4). For SSP1-RCP2.6, the species richness in this region increases mostly due to increases in forested land, non-forested land, and cropland, while only pasture substantially decreases in this region (Fig. 4). This is directly reflected in the relative loss of species richness of -1.4% +/-1.3% SD from climate change and -2% from land use change for pasture in West, Central, East and South Africa. While climate change and land use change generally have reinforcing negative or positive impacts, their impacts can also diverge, even compensating for each other (e.g., Oceania for forested land SSP1-RCP2.6 and SSP4-RCP6.0; Fig. 3).

**Fig. 4.**
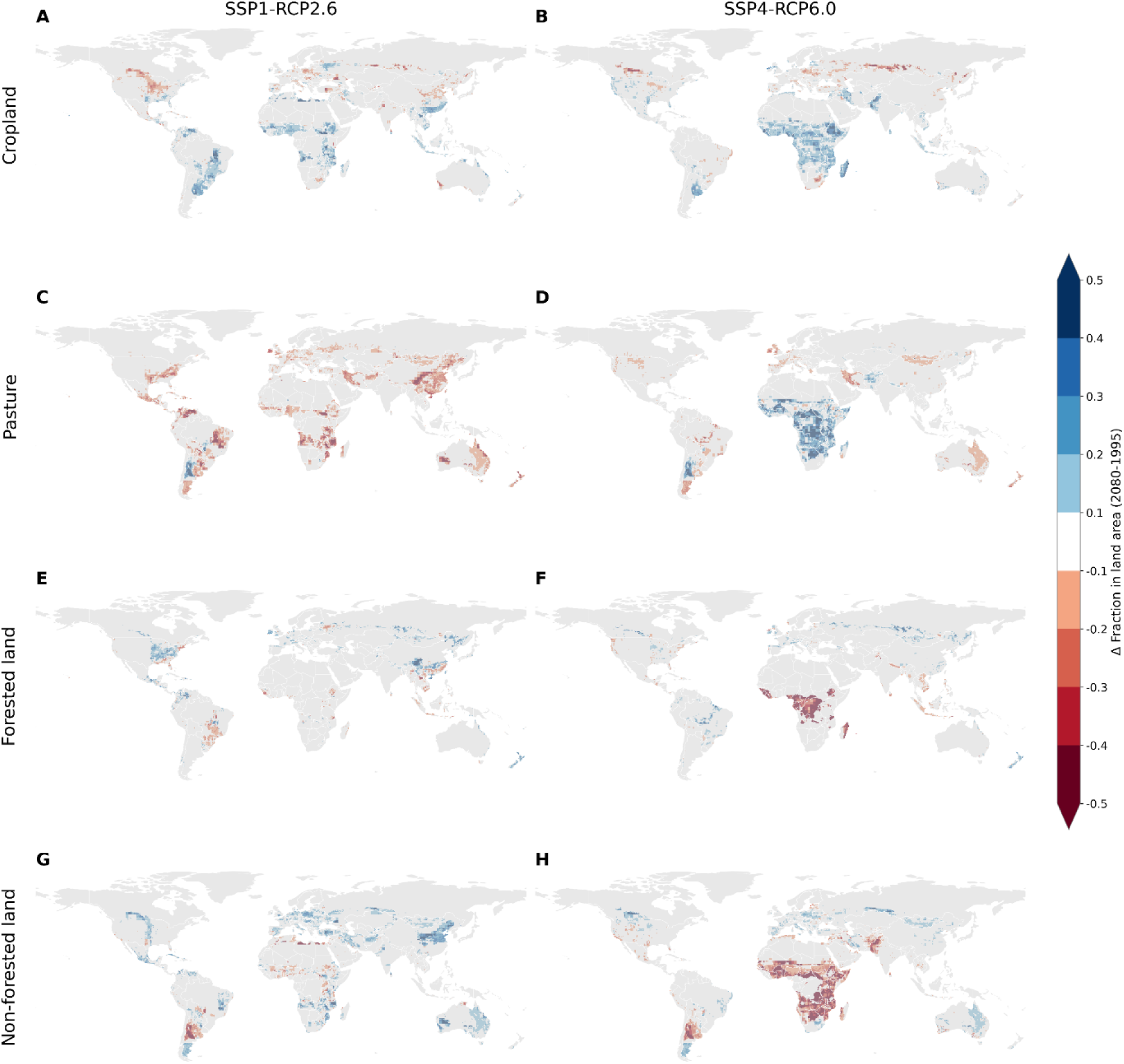
Projected change in land use category as fraction per grid cell for 2080 compared to 1995 based on LUH2 (Hurtt et al., 2020) for different land use categories for SSP1-RCP2.6 (A, C, E, G) and for SSP4-RCP6.0 (B, D, F, H). The changes are shown for four land use categories: cropland (A and B), pasture (C and D), forested land (E and F), and non-forested land (G and H). Blue indicates a decrease and red indicates an increase in land use as a fraction of the grid cell.

Eastern Europe and North America show the most notable relative gains in projected species richness over all land use categories and scenarios (Fig. 3). The majority of gain is climate-driven with the most pronounced species richness increase for the land use category ‘pasture’, where under SSP4-RCP6.0 22.6% +/-3.8% SD of the gain is due to climate change and 1.1% due to land use change (Fig. 3). This increase in the higher latitudes, however, is sensitive to the underlying dispersal assumption. Our additional sensitivity analysis with a no-dispersal assumption, where the future ranges correspond to the present ranges of the species, indicates less gain in the higher latitudes and more loss of species richness in the mid-latitudes (Fig. S3). When also including species with habitat suitability marked as “marginal” as part of the sensitivity analysis, the geographical patterns are consistent and there are only minor differences in the relative species richness change (Fig. S9).

### 3.3 Land area affected by species richness change

To put biodiversity impacts into perspective, we calculate the proportion of global land area affected by species richness gains and losses. In both scenarios and across all taxa, losses dominate under climate change only, and land area with losses being more widespread under SSP4-RCP6.0 (Fig. 5). 54.2% of land area loses species richness for mammals due to climate change in SS1-RCP2.6, while 37.4% of land area gains mammal species (Fig. 5C). For SSP4-RCP6.0, land area with species richness loss increases to 56.4%, and the proportion of area with gains also increases slightly to 37.6%, at the expense of the area with no change (Fig. 5I). The effect of land use change shows a more contrasted picture with a relatively equal proportion of losses and gains under SSP1-RCP2.6 (and even a predominance of gains for mammals with 48.5% of the land area) and a predominance of losses for the SSP4-RCP6.0 scenario (Fig. 5F & L). Also, for the other taxa the proportion of area leading to species richness loss from land use change is exacerbated for SSP4-RCP6.0 compared to SSP1-RCP2.6. The proportion of losses increases by 9.5% for birds (Fig. 5E & K) and by 6.5% for amphibians (Fig. 5D & J). It shows that land use change can be either a global alleviating (SSP1-RCP2.6) or exacerbating (SSP4-RCP6.0) factor of climate-induced biodiversity loss, although it does contribute to both regional losses and gains in both scenarios.

**Fig. 5.**
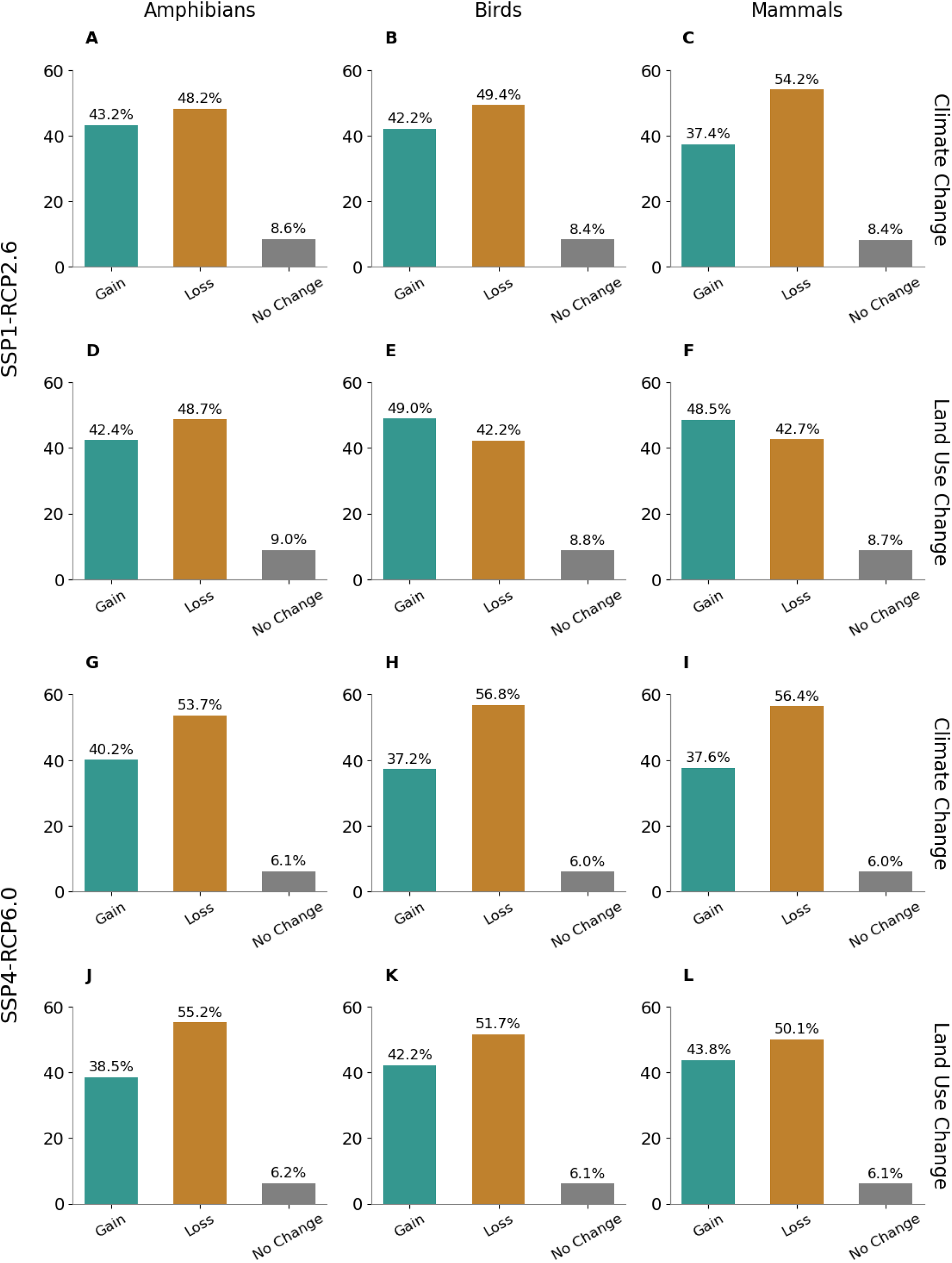
Climate and land use-driven changes in land area. Proportion of the total land area (excluding Antarctica) affected by species richness gain or loss for each taxon and scenario for climate and land use change. If the change equals 0, it was classified as no change. The species richness was first calculated as the summed probabilities of occurrence over all species for each taxon and scenario. For each combination of GCM and SDM, the probabilities are aggregated by calculating the latitudinal weighted mean. All future projections are then compared with the data of 1995. Based on this, the gains and losses are separated, and based on the grid cell we calculate the proportion of global area affected.

The proportion of losses is larger for all taxa under SSP4-RCP6.0 compared to SSP1-RCP2.6 for both climate and land use change. For mammals and birds, the proportion of global area with a climate-driven species richness loss is larger for SSP4-RCP6.0, with a loss of 56.4% and 56.8%, respectively, in comparison to the land use-driven species richness loss (50.1% and 51.7%, respectively). For amphibians, however, the projected share of global area with a species richness loss due to land use change is 55.2% and 53.7% from climate change.

## 4. Discussion

Our analysis has demonstrated that including land use change in future biodiversity projections leads to different species richness impacts than if investigating climate change impacts alone. While the climate change impact mostly scales with the climate scenario, the land use impact has a strong, more intricate scenario dependency. Indeed, we find that land use change can be either a global alleviating (SSP1-RCP2.6) or an exacerbating (SSP4-RCP6.0) factor of climate-induced biodiversity loss. Overall, our findings are in line with existing literature showing that climate change leads to a projected species shift towards higher latitudes, whereas land use change may limit this projected expansion (Methorst et al., 2017). Newbold (2018) also showed that the impacts of climate change and land-use change on biodiversity are expected to vary significantly across different regions. They highlight how these pressures are likely to combine and intensify in tropical grasslands, savannas, and the edges of tropical forests, leading to significant losses of species. In contrast, the heartlands of tropical forests and northern boreal regions might see less impact from land use changes but are still at risk of considerable biodiversity loss due to climate change. Meanwhile, temperate regions could experience relatively minor changes in species richness as a result of future climate change (Newbold, 2018).

The increase in forested and non-forested land over the course of the century projected for SSP1-RCP2.6 directly translates into species richness change by habitat and into overall species richness changes as most of the species included in our study have either forested (78.5%) or non-forested land (65.2%) or both categories as suitable habitats (Table 1, Fig. 3). Especially, West, Central, East and South Africa shows that under increasing forest and non-forested land, species richness in 2080 is projected to increase compared to 1995 levels and land use change will be the main driver of this change. However, this positive effect strongly depends on the quality of the land use change. For example, whether forest gains benefit biodiversity will depend on the type of reforestation. In the tropics, most reforestation currently occurs with non-native species and afforestation on natural land with high biodiversity, such as savannas, which can also be detrimental to biodiversity (Parr et al., 2014).

Furthermore, the land use scenarios used for this study, especially the SSP1-RCP2.6 scenario, assume a biodiversity-conscious world (Hurtt et al., 2020). Projections from other IAMs under the same scenario have been shown to result in more negative outcomes for biodiversity (Leclère et al., 2020). Many other IAM projections under this scenario assume a more bioenergy-extensive world (Hof et al., 2018), whereas our LUH2 projections driven by GCAM and IMAGE assume reforestation and afforestation in high-and medium-income countries with an assumption of global forest cover increase (Hurtt et al., 2020). Thus, even resulting in an increase in forested land in South America under SSP4-RCP6.0 (Fig. 4).

Our results show that when forested and non-forested land are projected to decrease under SSP4-RCP6.0, the species richness loss amounts to more than 30% compared to 1995 values in West, Central, East and South Africa. Other studies have also projected similar declines in sub-Saharan African species richness (Visconti et al., 2016) because increases in human population led to a reduction in natural vegetation (PBL, 2012). Our results have shown that only regions in the higher latitudes show an increase in species richness in the future, even for the SSP4-RCP6.0 scenario (Fig. 3).

Many studies have discussed species’ range shifts towards higher latitudes as a response to climate change (Chen et al., 2011; Lenoir & Svenning, 2015; Parmesan & Yohe, 2003; Ramalho et al., 2023). Our results show a more complex pattern for land use change impacts on biodiversity but are in line with other studies that evaluate habitat suitability and risks, such as the study by Powers and Jetz (2019), which underscores significant global habitat declines and the urgency of national stewardship in biodiversity-rich regions like South America, Southeast Asia, and Africa. Their work also highlights the critical need to identify and prioritize the most vulnerable species and locations, a point that aligns with our findings and reinforces the importance of proactive conservation planning in mitigating biodiversity threats (Powers & Jetz, 2019). The species richness increase in the higher latitudes can be attributed to climate change and highly depends on the underlying dispersal assumption. Interestingly, the mid-latitudes are projected to profit from land use change in the future. Our results suggest that countries known to create a large part of their biodiversity footprints outside their own country’s borders by importing large amounts of agricultural products from regions with high biodiversity loss (Schwarzmueller & Kastner, 2022), namely, countries in Western Europe, North America, and the Middle East show a projected increase in future land use change-driven species richness. This has not only implications for conservation planning but also for global equity discussions regarding the externalization of biodiversity loss at the benefit of national biodiversity protection.

Our results were computed under a limited-dispersal assumption. However, a sensitivity analysis using a no-dispersal scenario showed that they are still optimistic (Fig. S3). The underlying dispersal ability assumptions are important to consider when analyzing climate change impacts on species, as different assumptions can have substantial effects on the projected future ranges (Methorst et al., 2017; Newbold, 2018; Thuiller et al., 2019). In addition, we have tested the sensitivity of the results towards our habitat suitability assumptions. We performed an additional analysis including species with “marginal” suitability according to the IUCN classification alongside the “suitable” habitat. This inclusion resulted in minimal changes in the resulting patterns of projected species richness change (Fig. S9). But despite this little sensitivity of the results towards our habitat suitability assumptions, it is important to note that our analysis includes only four land use categories, which is a crude summary of 104 categories that are available in the IUCN Habitat Classification Scheme. The summary of primary forested and secondary forested land further misses reforestation and afforestation activities. This could potentially confound our results, as it assumes that a species that has lost its primary forest can simply find habitat in a secondary forest. Although evidence from the Amazon shows that at least some species groups are not faring well in secondary forests or plantations (Barlow et al., 2007). Furthermore, non-forested land including for example both savannahs and wetlands are harboring very specific species closely tied to their habitats, which in our study are not separated. The assumption to combine secondary and primary forest due to their potential complementarity might also present disadvantages, as it does not fully account for species that are closely tied to primary forest habitat. Also, we lack “urban” as a land use category, which is an important habitat for many bird species (Ortega-Álvarez & MacGregor-Fors, 2009). This is especially relevant to our study since more than half of the species considered in this study are birds. Going forward, it would thus be important to include the impacts of urban areas and urbanization in a combined climate and land use change study. However, integrating urban areas into such studies requires specific information about species for which “urban” areas serve as suitable habitats – information currently not part of the IUCN habitat classification scheme. But a study by Simkin et al. (2022) highlights the rapid urbanization expected in biodiversity-rich regions, including sub-Saharan Africa, South America, Mesoamerica, and Southeast Asia and underscores the urgent need to consider urban land in conservation strategies. Despite these simplifications we assume our categories to be constructive for a global analysis, which is key to understanding and identifying global biodiversity patterns. An important area for future discussion is the combined effects of climate and land-use change on biodiversity. While this study investigates the separate and combined impacts of these two drivers, it is crucial to acknowledge that they often interact synergistically, potentially leading to even greater biodiversity loss than the sum of their individual effects. Additionally, this analysis focuses on just two of many anthropogenic drivers (Jaureguiberry et al., 2022). As Rillig et al., (2019) highlight, including more factors like pollution, invasive species, and habitat fragmentation can significantly alter the projected outcomes. A broader consideration of these combined and interacting pressures will provide a more realistic picture of future biodiversity change.

Besides the importance of climate and land use change impacts on species ranges as well as the significance of the choice of scenarios and their underlying narratives, we have also quantified the uncertainty from the set of two SDMs (GAM and GBM) and four GCMs (MIROC5, GFDL-ESM2M, HadGEM2-ES, and IPSL-CM5A-LR) used in this study. However, we could not assess structural uncertainty from the land use projections, as each scenario in our analysis is derived from a single IAM, based on the SSP marker framework. The selection of marker scenarios -e.g., IMAGE for SSP1 and GCAM for SSP4-is part of a standardized and deliberate process within the SSP framework. Marker scenarios were chosen based on each IAM’s ability to represent the core characteristics of a given SSP narrative and to ensure internal consistency and interpretability across the scenario space (Riahi et al., 2017). This approach provides a coherent comparison across contrasting futures. However, it also means that differences in outcomes may partially reflect structural differences between IAMs. Other IAMs have implemented the same SSPs, and future research should incorporate multiple IAMs per scenario to better quantify and address this source of uncertainty (e.g. Molina Bacca et al., 2024). Future studies should focus on including a wider range of IAMs especially since we have shown that the contribution from land use change is substantial and tied to climate change impacts. Additionally, opportunities for further improvement and refinement regarding the use of SDMs need to be recognized. First, the SDMs were run using expert range maps from IUCN (IUCN, 2023) and Birdlife International and Nature Serve (Birdlife International, 2015). The extent of occurrence expert range maps tends to overestimate the actual species distribution (Piirainen et al., 2023) and can be imprecise or incomplete (IUCN, 2023). Second, only species are included in the study that could properly be modeled by SDMs, species with a range smaller than ten grid cells were excluded from the SDM analysis as otherwise, the combination of few occurrences and many predictor variables can easily lead to model overfitting (Breiner et al., 2015; Hof et al., 2018; Platts et al., 2014). Consequently, many narrow-ranging species that are most threatened to go extinct are disregarded from our study. We recognize that this method may be unsuitable to draw conclusions about the impacts of climate and land use change on individual species. Still the method does allow inferring general patterns across a wide array of species.

Overall, we have shown that species richness losses predominate in terms of affected global land area for all scenarios and taxa under the combined effect of climate and land use change. This underlines the importance of climate mitigation efforts to curb the rise in global mean temperature. Furthermore, when we recalculate the loss in species richness to reflect land use change specifically, we found a greater projected loss under SSP4-RCP6.0 scenario compared to SSP1-RCP2.6 for all three taxonomic groups. Thus, land use policies will have a key role in fighting global biodiversity loss, especially in lower latitudes. Conservation efforts should therefore account for both drivers, climate and land use change, according to their respective impact. Finally, the regional differences shown in this study highlight the crucial need for region-specific conservation strategies.

## Competing interests

The authors declare that they have no competing interests.

## Data and Code Availability Statement

The data generated and analyzed during the current study are available from the corresponding author. The data underlying the main analysis and figures are uploaded to Zenodo and are available from https://doi.org/10.5281/zenodo.11119306. All code used in this study is available from https://doi.org/10.5281/zenodo.14093900.

## Notes

https://doi.org/10.5281/zenodo.11119306

https://doi.org/10.5281/zenodo.14093900

### Competing Interest Statement

The authors have declared no competing interest.

### Summary of Updates

This version of the manuscript has been revised.

