## Supplementary for "Future climate and land use change will equally impact global terrestrial vertebrate diversity"

**This PDF file includes:**

Supplementary Text
Figs. S1 to S9
Table S1

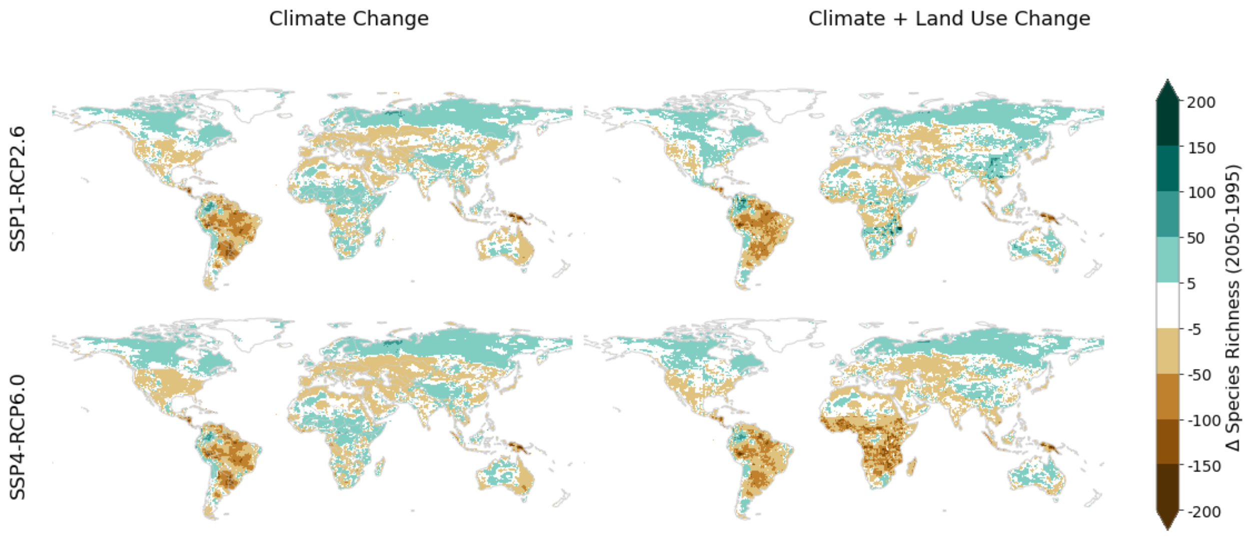


**Fig. S1.**

Projected species richness for the year 2050 compared to 1995 under climate change only with the 1995 land use as a baseline for the present and future and climate and land use change with present and future land use data respectively. Results are shown for SSP1-RCP2.6 and SSP4-RCP6.0. Species richness is calculated as the summed probabilities of occurrence over all species of the taxa amphibians, birds, and mammals. The mean over all GCM and SDM combinations is shown here.

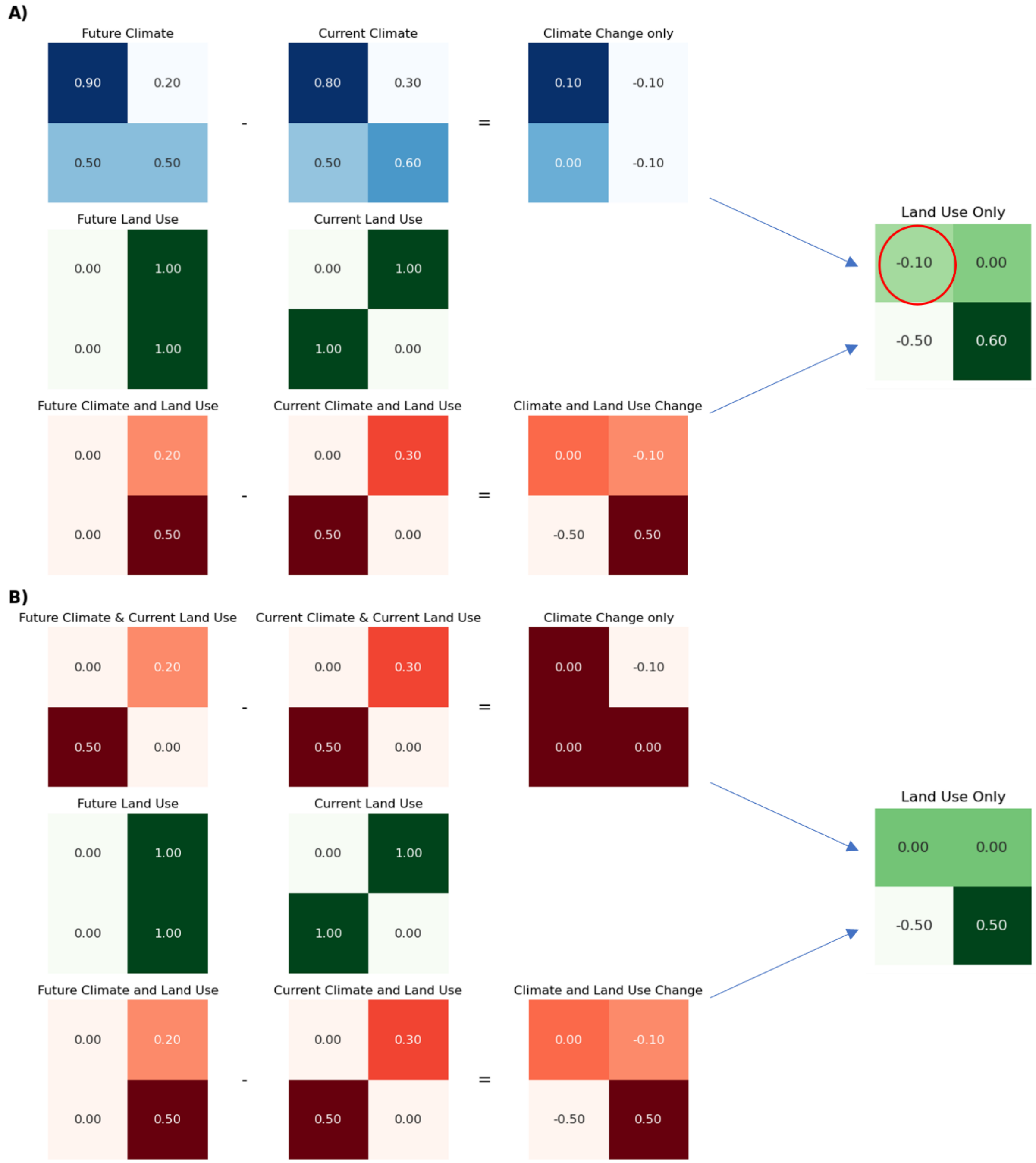


**Fig. S2.**

Schematic overview of the different methodological steps for the calculation of the land use residual based on the initial method calculating the land use-only change as a residual directly from the probabilities of occurrences (i.e. the SDM output unfiltered) (**A**) and in this study used method, where climate change impact is filtered by the 1995 land use as a baseline and then climate- only is calculated as the residual of the future and present with constant land use change (**B**).

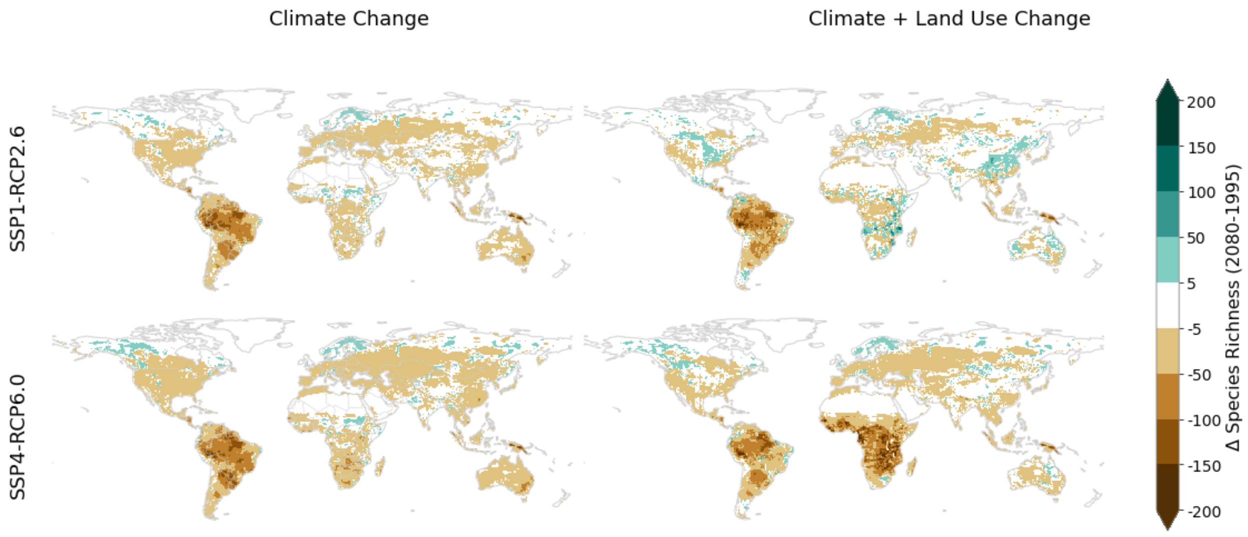


**Fig. S3.**

Projected species richness for the year 2080 compared to 1995 under climate change only with the 1995 land use as a baseline for the present and future and climate and land use change with present and future land use data respectively for a no dispersal assumption. Results are shown for SSP1-RCP2.6 and SSP4-RCP6.0. Species richness is calculated as the summed probabilities of occurrence over all species of the taxa amphibians. The mean over all GCM and SDM combinations is shown here.

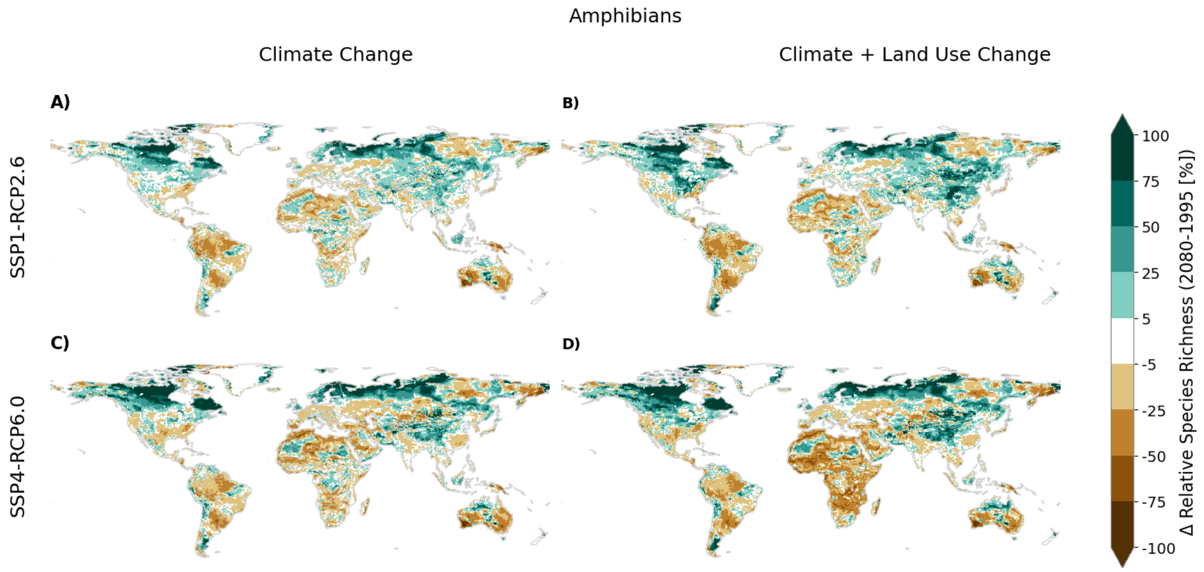


**Fig. S4.**

Projected species richness for the year 2080 compared to 1995 under climate change only with the 1995 land use as a baseline for the present and future and climate and land use change with present and future land use data respectively. Results are shown for SSP1-RCP2.6 and SSP4-RCP6.0. Species richness is calculated as the summed probabilities of occurrence over all species of the taxa amphibians. The mean over all GCM and SDM combinations is shown here.

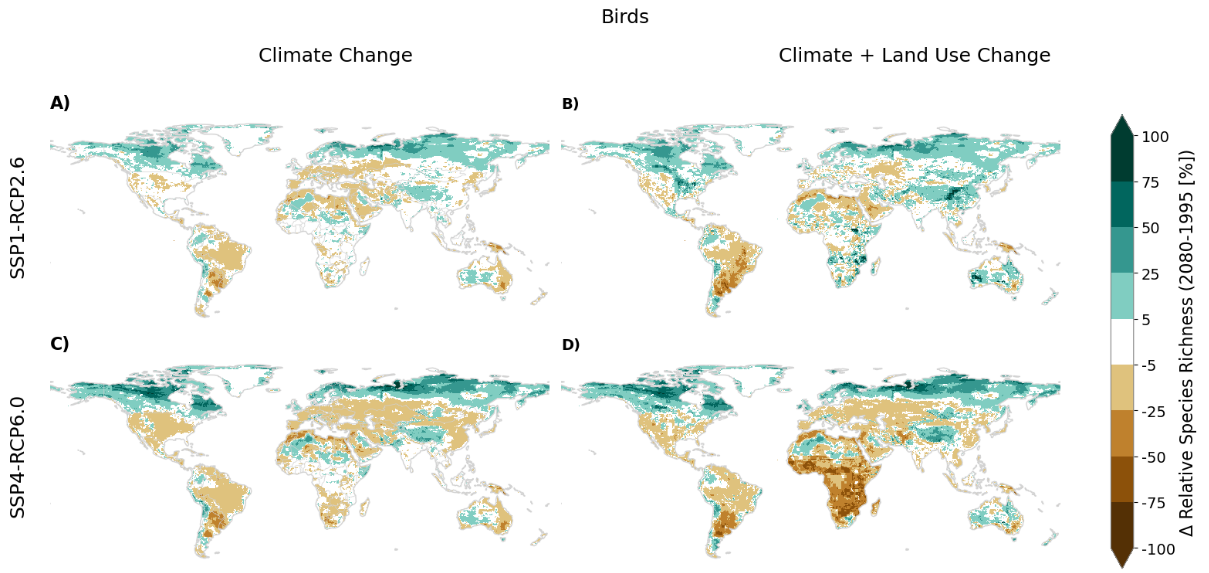


**Fig. S5.**
Projected species richness for the year 2080 compared to 1995 under climate change only with the 1995 land use as a baseline for the present and future and climate and land use change with present and future land use data respectively. Results are shown for SSP1-RCP2.6 and SSP4-RCP6.0. Species richness is calculated as the summed probabilities of occurrence over all species of the taxa birds. The mean over all GCM and SDM combinations is shown here.

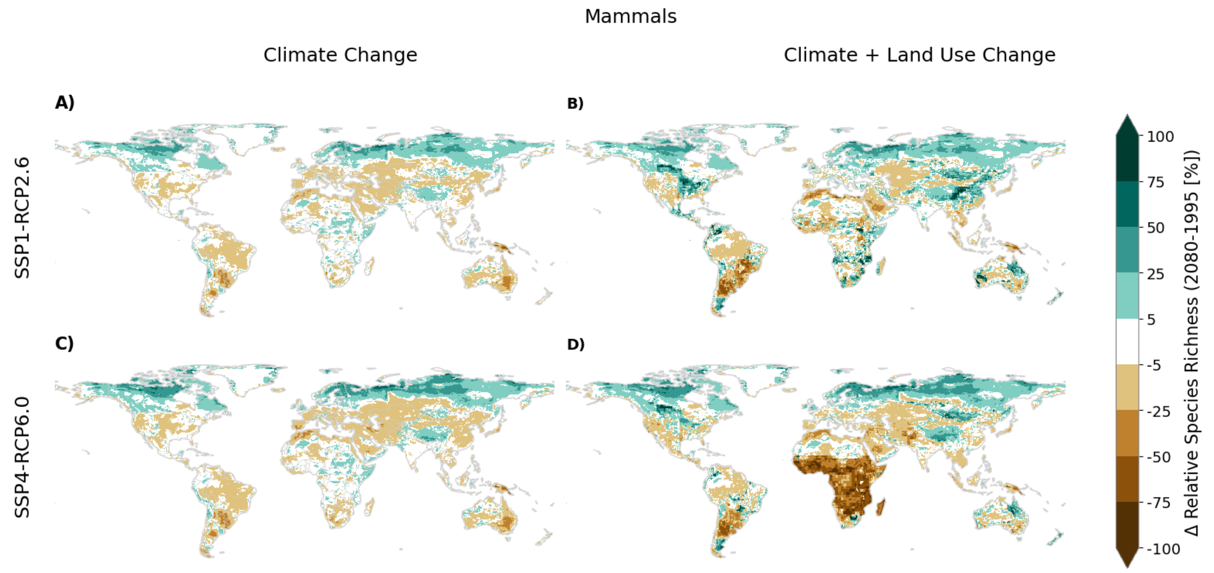


**Fig. S6.**

Projected species richness for the year 2080 compared to 1995 under climate change only with the 1995 land use as a baseline for the present and future and climate and land use change with present and future land use data respectively. Results are shown for SSP1-RCP2.6 and SSP4-RCP6.0. Species richness is calculated as the summed probabilities of occurrence over all species of the taxa mammals. The mean over all GCM and SDM combinations is shown here.

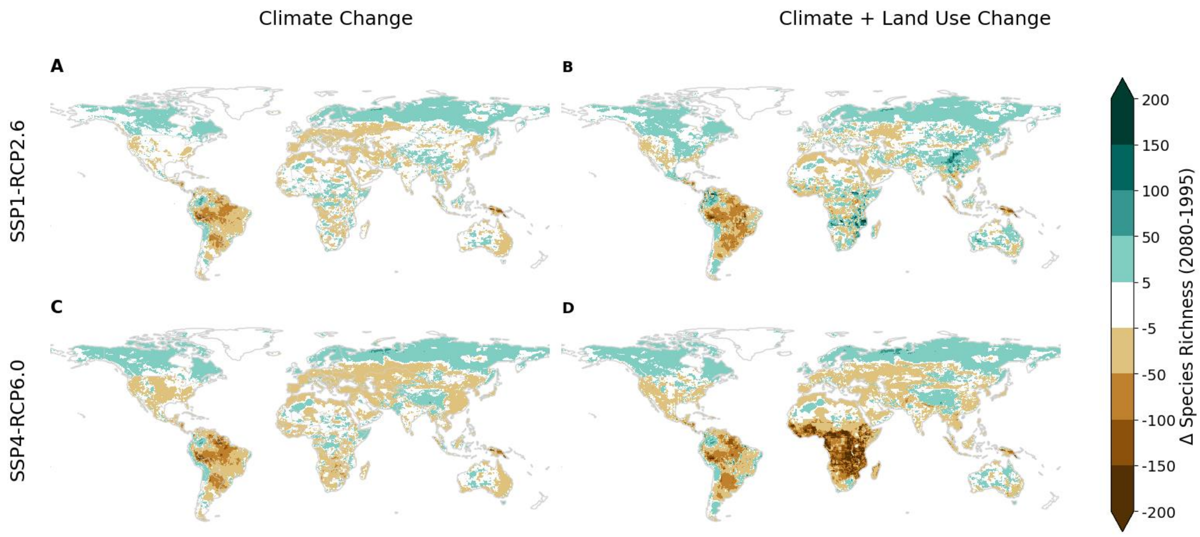


**Fig. S7.**

Projected absolute species richness for the year 2080 compared to 1995 under climate change only with the 1995 land use as a baseline for the present and future and climate and land use change with present and future land use data respectively. Results are shown for SSP1-RCP2.6 and SSP4-RCP6.0. Species richness is calculated as the summed probabilities of occurrence over all species of the taxa amphibians, birds, and mammals. The mean over all GCM and SDM combinations is shown here.

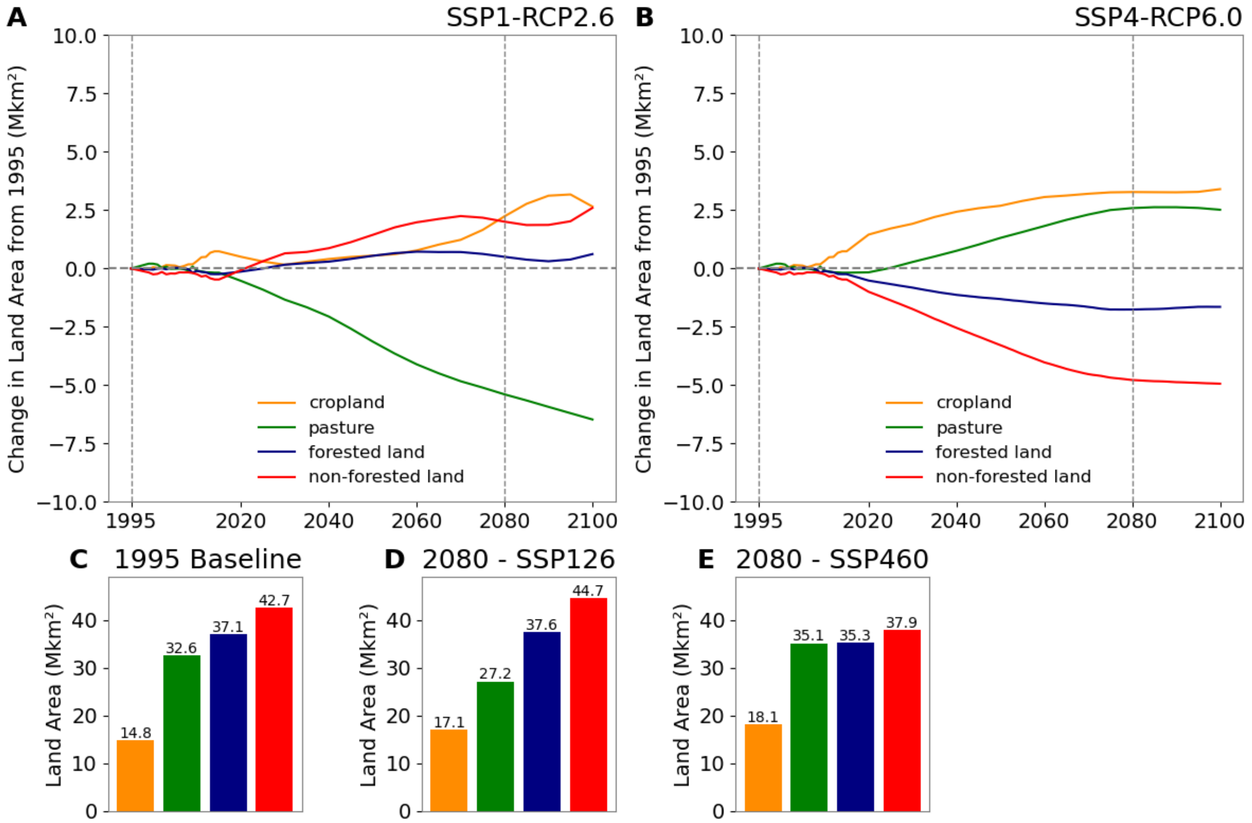


**Fig. S8.**

Future land use scenarios. Land use change in (A) the “sustainability” (SSP1-RCP2.6) from IMAGE (24; see Material and Methods) scenario and (B) the “inequality” (SSP4-RCP6.0) scenario from GCAM (25; see Material and Methods), in reference to the year 1995 based on the LUH2 dataset (22) as well as (C) the land area per land use category for 1995, (D) for 2080 under SSP1-RCP2-6) and (E) for 2080 under SSP4-RCP6.0).

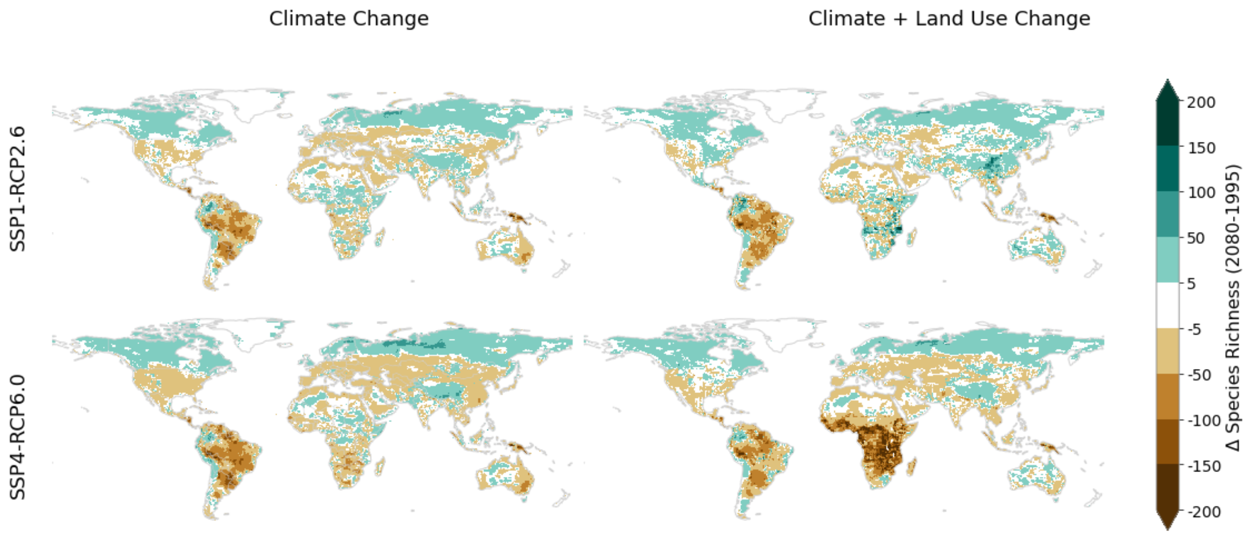


**Fig. S9.**

Projected species richness for the year 2080 compared to 1995 under climate change only with the 1995 land use as a baseline for the present and future and climate and land use change with present and future land use data respectively. All habitats labeled as "suitable" and "marginal" by the IUCN Habitat Classification Scheme are included. Results are shown for SSP1-RCP2.6 and SSP4-RCP6.0. Species richness is calculated as the summed probabilities of occurrence over all species of the taxa amphibians. The mean over all GCM and SDM combinations is shown here.

**Table S1.**

Regions used in this study and their connection to the IPBES subregions (IPBES, 2019).

| **Regions in this study** | **IPBES subregions** |
|---|---|
| Caribbean & Mesoamerica | Caribbean |
| | Mesoamerica |
| West, Central, East & South Africa | Central Africa |
| | East Africa and adjacent islands |
| | Southern Africa |
| | West Africa |
| Central and Western Europe | Central and Western Europe |
| Central, North-East & South Asia | Central Asia |
| | Nort-East Asia |
| | South Asia |
| Eastern Europe | Eastern Europe |
| North Africa & Western Asia | North Africa |
| | Western Asia |
| North America | North America |
| Oceania | Oceania |
| South America | South America |
| South-East Asia | South-East Asia |
